# The effectiveness and cost effectiveness of a hospital avoidance program in a residential aged care facility

**DOI:** 10.1101/523969

**Authors:** Hannah Carter, Xing Lee, Trudy Dwyer, Dee Jeffrey, Barbara O’Neill, Chris Doran, Lynne Parkinson, Sonya Osborne, Kerry Reid-Searl, Nicholas Graves

**Affiliations:** Australian Centre for Health Services Innovation, Institute of Health and Biomedical Innovation, Queensland University of Technology; Central Queensland University; PresCare; Central Queensland University, University of Connecticut; Centre for Indigenous Health Equity Research, Central Queensland University

**Keywords:** Cost-effectiveness, residential age care facility, hospital avoidance, nursing home, hospital admission

## Abstract

**Background:** Residential aged care facility residents experience high rates of hospital admissions which are stressful, costly and often preventable.

**Design:** Prospective pre-post cohort study and decision model analysis

**Intervention:** A decision-support tool was implemented to enable nursing staff to detect, refer and quickly respond to early signals of a deteriorating resident. Advanced clinical skills training, new diagnostic equipment and guided support from clinical lead nurses and nurse practitioners was provided to support nursing staff in the delivery of appropriate sub-acute care.

**Outcome measures:** Rate of hospital admissions; length of stay; incremental cost per QALY; net monetary benefit.

**Results:** The hospital avoidance program was associated with a 19% reduction in annual hospital admissions and a 31% reduction in the average length of stay. When modelled in a cohort of 1,000 residents the program resulted in a total of 1,606 fewer hospital bed days per annum. This contributed to a total cost saving of $2.6 million and 0.62 incremental QALYs gained per 1,000 residents. The program had a positive net monetary benefit and was considered cost-effective, even when the willingness to pay for health care gains was set to zero. A probabilistic sensitivity analysis estimated that there was an 86% probability that the program was cost-effective after taking the uncertainty of the model inputs into account.

**Conclusions:** This study provides compelling evidence for the effectiveness and cost-effectiveness of a RACF nurse led sub-acute care program in preventing unnecessary hospital admissions.

## Introduction

Ageing populations have contributed to growing demand for aged care services internationally.^1-4^ In Australia, admissions to aged care services have increased by 31% over the last decade ^5^. It is known that residents of residential aged care facilities (RACF) are frequent users of hospital services, with annual rates of more than 30 hospital transfers per 100 RACF beds commonly reported ^6^. These admissions account for 3% of all hospital bed days ^7^.

Hospital admissions in this cohort are costly, with the average cost per an admitted RACF resident estimated at $1,028 per bed day in a 2011 Australian study^8^. Admissions are considered stressful and are often unnecessary or potentially preventable.^9-14^ Residents and their families express a preference for care to be provided in their home ^15^, and older people treated in these settings are less likely to experience complications commonly incurred during hospitalisation ^6 16-18^. Previous studies have found that RACF nursing staff have a genuine desire to care for their acutely unwell residents within the facility^19-22^. There is therefore a strong clinical and economic basis for hospital avoidance interventions that promote appropriate nursing care within the RACF.

Evidence is emerging that hospital admissions from the RACF can be reduced by implementing models of care that improve nursing staff confidence, clinical skills and access to resources ^6 23-28^ Previous studies have focussed on the impact of these programs on emergency department (ED) transfers and hospital admissions, with few reporting on changes to average length of stay for admitted patients. There is no published evidence on the cost-effectiveness of making these changes to models of care.

The Early Detection of Deterioration In Elderly residents (EDDIE) program is a hospital avoidance intervention aimed at improving the proactive care and management of residents by RACF nursing staff. The objectives of this study were to estimate the impact of the EDDIE intervention on hospital admission rates and length of stay; and, report on the cost-effectiveness of the EDDIE intervention as compared to usual care.

## Methods

A prospective pre-post cohort study design was adopted to estimate the changes to hospital admission rates and length of stay in the 12 months pre and post-implementation of the EDDIE intervention in a 96 bed regional Australian RACF in June 2016. Participants included all residents within the facility over the study period. This represented a range of 91 to 96 residents, with an average monthly occupancy of 94 residents observed across both the pre and post EDDIE cohorts. We refer to residents present during the 12 months post implementation of the EDDIE intervention as the intervention cohort (June 2016 – May 2017), and residents present during the 12 months prior to the EDDIE interventions as the usual care cohort (June 2015 - May 2016). The study was approved by the relevant Human Research Ethics Committee (reference number: CQU/H14/01-012). We used the CHEERS checklist as our reporting guide ^29^.

### The EDDIE Intervention

The intervention involved the use of a “traffic light” decision support tool which assisted nursing staff to detect, refer and quickly respond to early signals of a deteriorating resident. The program included advanced clinical skills training, new diagnostic equipment and guided support from clinical lead nurses and nurse practitioners. Critically, the intervention did not involve the employment of additional nursing staff within the RACF. The focus was instead on upskilling existing staff members and empowering them to manage sub-acute episodes within the facility which required intensive treatments, interventions and frequent assessments.

### Statistical analysis of the observed data

The impact on variation in the data was explored by fitting statistical distributions around key results based on the observed means and standard deviations from both intervention and usual care cohorts. A normal distribution provided the best fit for the number of admissions per annum. A gamma distribution was used to represent length of stay as its positive, right-skewed nature accounted for a small proportion of admissions experiencing relatively long lengths of stay. Probability density functions were then produced by randomly drawing 1,000 samples from the respective distributions.

### Costs of implementation

A set of the initial implementation costs of EDDIE were estimated based on the project data collection. The decision support tool was developed and piloted in a previous study and the costs associated with this were not included in this analysis. We accounted for the cost of printing the decision support materials, as well as the staff costs associated with the implementation strategy such as training, stakeholder engagement and project management activities. The costs of staff time were assigned using published salary band data where available. These costs are reported in Appendix 1. Due to the one-off, upfront nature of these costs they were not included in the modelled analysis.

### Modelled cost-effectiveness analysis

A Markov model was developed to estimate the cost-effectiveness of the EDDIE intervention compared to usual care over a period of 12 months in a cohort of 1,000 residents. The model defined a number of discrete health states that aged care residents could experience over a period of 365 days including time spent within the RACF as a typical resident, sub-acute episodes managed within the RACF, hospital admissions and death. A set of transition probabilities governed the likelihood of residents transitioning from one state to another at the end of each daily cycle. The Markov process and health state transitions for both the EDDIE intervention and usual care are represented in Figure 1.

**Figure 1.**
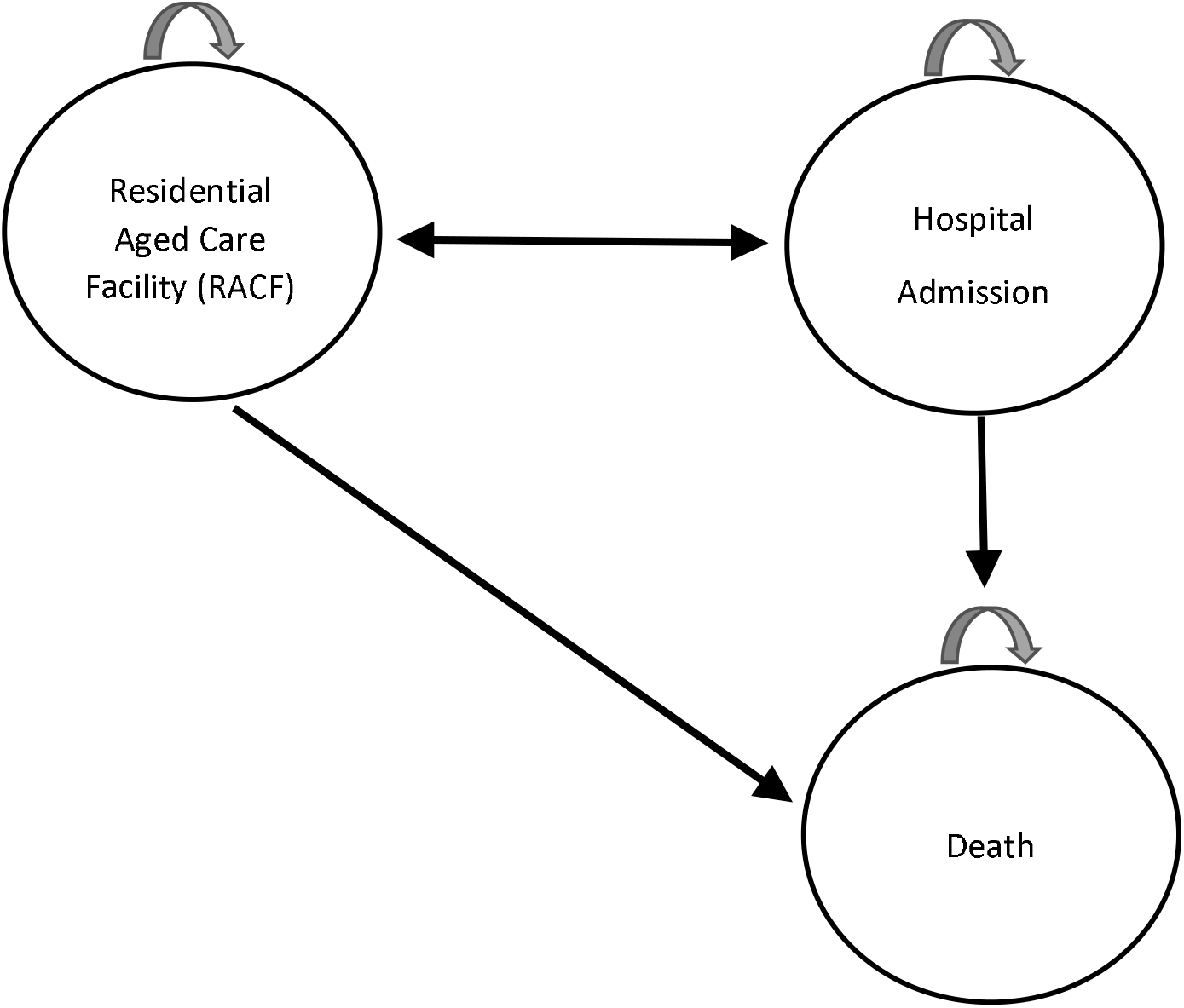
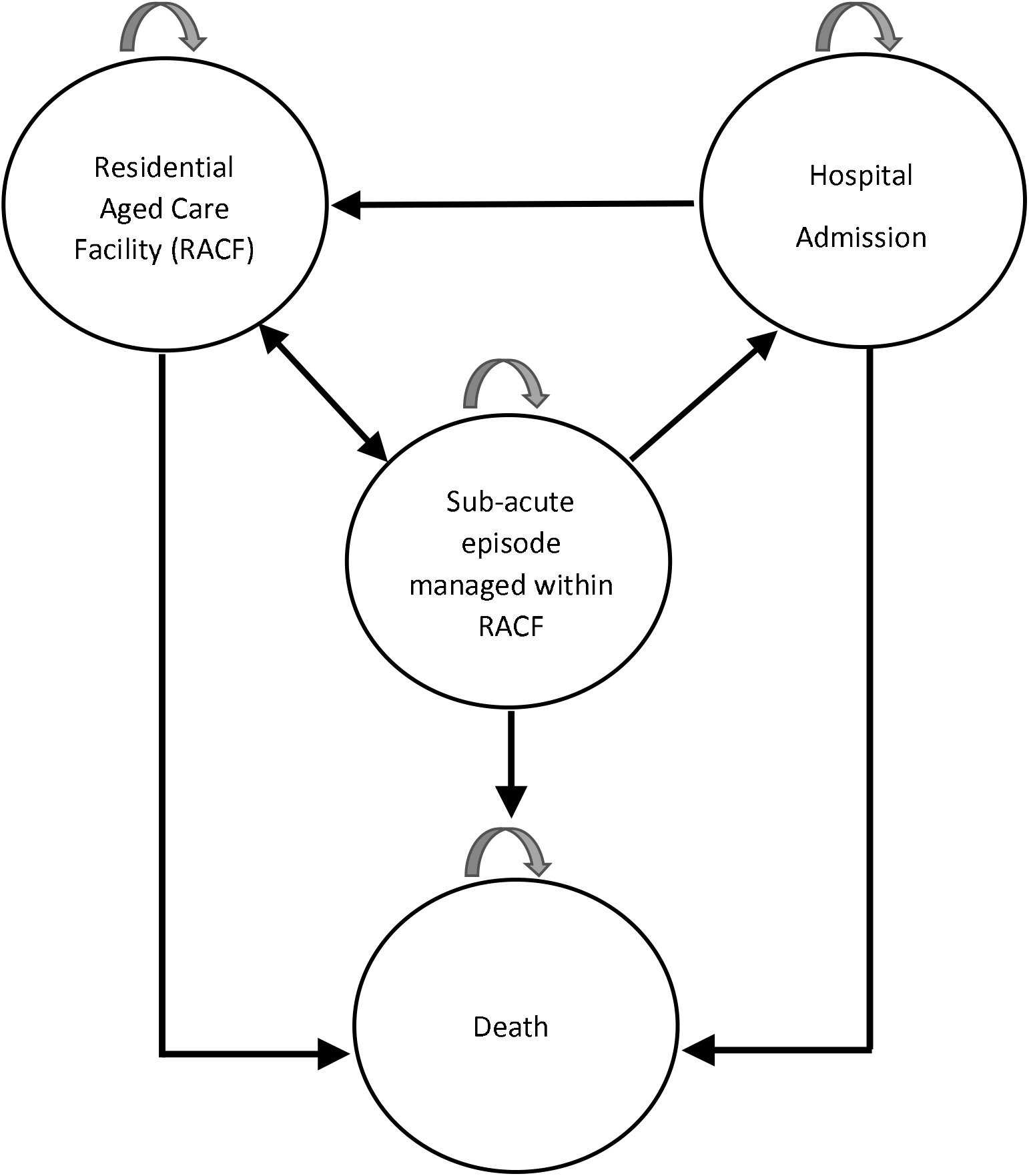
Markov model structure. **A: Health states and transitions within usual care:** All residents begin the model in the residential aged care facility (RACF). Residents who deteriorate are admitted to hospital for treatment, and return to the RACF once stable. Residents may die in the RACF or in hospital. **B: Health states and transitions within the EDDIE intervention:** All residents begin the model as stable in RACF. Residents who deteriorate are first managed as a sub-acute episode within the RACF, where they may then stabilise and remain within the RACF. Residents who deteriorate further are admitted to hospital for treatment, and return to the RACF once stable. Residents may die at any point in the model.

The model was used to synthesise data collected in the study with published literature on the outcomes associated with relevant health states experienced by residents. Incremental cost-effectiveness was measured in terms of the cost per quality adjusted life year (QALY) gained. QALYs were derived by weighting the time spent in each health state by a health related quality of life value (utility) associated with that state. A utility of zero is equivalent to death and a utility of 1 is equivalent to full health ^30^. The evaluation was conducted from the perspective of the Australian health care system in which aged care services and hospital admissions are publicly funded. All costs are reported in 2018 Australian dollars.

### Model inputs

All probabilities, costs and utility values applied in the model, along with respective standard deviations and data sources where relevant, are reported in Table 1.

**Table 1.** Transition probabilities applied in the cost-effectiveness model.

| Parameters | Base case estimate | SD | Source |
| --- | --- | --- | --- |
| **Transition probabilities:** |  |  |  |
| Intervention cohort |  |  |  |
| Daily probability of sub-acute episode | 0.003 | 0.007 | Study data |
| Proportion of sub-acute episodes treated within the facility | 0.670 | 0.388 | Study data |
| Daily probability of sub-acute episodes admitted to hospital | 0.722 | 0.288 | Study data |
| Daily probability of residents being discharged from hospital | 0.283 | 0.150 | Study data |
| Usual care cohort |  |  |  |
| Daily probability of residents being admitted to hospital | 0.001 | 0.004 | Study data |
| Daily probability of residents being discharged from hospital | 0.151 | 0.072 | Study data |
| All residents |  |  |  |
| Daily probability of death | 0.0011 | 0.0001 | Study data |
| **Costs** |  |  |  |
| New diagnostic equipment (annualised)* |  |  |  |
| Bladder Scanner x1 | 1714 | 672 | Study data |
| ECG Machine x1 | 351 | 138 | Study data |
| Vital Signs Monitor x1 | 277 | 109 | Study data |
| RACF bed day | 194 | 76 | [31] |
| Ambulance transfer to hospital | 649 | 254 | [32] |
| Hospital bed day | 1807<sup>^</sup> | 708 | [8] |
| **Utility values** |  |  |  |
| RACF residents | 0.514 | 0.252 | [33] |
| Elderly inpatients admitted from RACF | 0.44 | 0.4 | [34] |

RACF = residential aged care facility; SD = standard deviation; ECG = electrocardiogram
\*Costs were annualised over a useful life of seven years according to Australian government depreciation schedules (Income Tax Assessment Act, Income Tax (Effective Life of Depreciating Assets) Determination 2015)
<sup>^</sup>Inflated to 2018 dollars using an index of hospital price inflation<sup>35</sup>

Transition probabilities were derived from the observed daily events data collected at the RACF over the period June 2015 - May 2016 for usual care and June 2016 - May 2017 for the EDDIE intervention.

Costing items included the cost of additional diagnostic equipment not typically utilised in the RACF setting that was purchased in order for trained staff to better detect and manage sub-acute episodes. Equipment costs were annualised over a period of seven years, reflecting their useful life as defined in the Australian government depreciation schedules^36^. A cost per day was assigned to RACF bed days based on current national fee schedules^31^. The cost of a hospital bed day was informed by a 2011 Australian study that produced estimates of admissions costs and length of stay that were specific to a RACF cohort^8^; this was then inflated to 2018 dollars using an index of hospital price inflation^35^. The cost of an ambulance transfer was also assigned with each hospital admission in line with standard practice^32^.

The model assigned separate utility values according to whether a resident was in the RACF or in hospital.

### Sensitivity analysis

A probabilistic sensitivity analysis was performed in order to estimate the impact of simultaneous uncertainty across all modelled estimates. A normal distribution was applied to cost parameters with a 95% confidence interval encompassing a variation of 20% above and below the base case estimate. The exception was the cost per hospital bed day which was assigned a gamma distribution (SD 1,028) based on the nature and availability of these data ^8^. Beta distributions were fitted to the transition probability and utility estimates using the standard deviations reported in Table 1. A Monte Carlo simulation was then performed with 1,000 randomly drawn samples taken from each of the modelled parameter distributions.

The modelled uncertainty was represented in the form of a distribution around the Net Monetary Benefit (NMB) associated with a decision to adopt the EDDIE intervention. This provides a measure of the value of the intervention in monetary terms when the willingness to pay for a QALY is known. A positive NMB indicates that an intervention is cost-effective. The NMB was estimated using a recently published study of the optimal willingness to pay for a QALY in an Australian setting of $28,000 ^37^. A sensitivity analysis estimated the cost-effectiveness of the intervention where the willingness to pay for health benefits was set to zero.

## Results

There were 112 sub-acute episodes recorded in the intervention cohort over 12 months, with 75 of these treated within the RACF only. The remaining 37 sub-acute events resulted in hospital admissions with a mean length of stay of 4.8 days. In comparison, a total of 45 hospital admissions over 12 months were recorded in the usual care cohort with a mean length of stay of 7.7 days. This represented a 19% reduction in annual hospital admissions and a 31% reduction in the average length of stay following implementation of the EDDIE intervention. Appendix 2 presents the probability density functions around both the admission rates and length of stay outcomes.

The modelled cost-effectiveness analysis estimated that the EDDIE intervention would prevent 95 hospital admissions in a cohort of 1,000 residents, resulting in a total of 1,606 fewer bed days (Table 2). This contributed to a total cost saving of $2.6 million per 1,000 residents treated. The incremental QALYs gained was positive, but modest at 0.62 QALYs per 1,000 residents. This was due to the relatively small decrement in utility associated with hospital admissions when compared to the baseline utility score of RACF residents.

**Table 2.** Mean cost-effectiveness outcomes taken from 1,000 Monte Carlo simulations modelled over 12 months in a cohort of 1000 residents.

| Modelled outcomes per 1,000 residents | Intervention | Usual care | Difference |
| --- | --- | --- | --- |
| Number of admissions | 270 | 365 | -95 |
| Total hospital bed days | 1,377 | 2,983 | -1,606 |
| Total costs (\$000's) | 61,889 | 64,483 | -2,594 |
| Total QALYs | 414.04 | 413.42 | 0.62 |
| Cost-effectiveness result | Intervention is dominant* |  |  |

\*A cost-effectiveness result of 'dominant' indicates an intervention is both more effective and less costly than the alternative.

The mean NMB of the EDDIE intervention over 1,000 Monte Carlo simulations was $2,611 per resident (SD $2,802) when adopting a willingness to pay of $28,000 per QALY ^37^. The distribution of NMB samples is presented in Appendix 3. Approximately 86% of the simulations produced a positive NMB, providing a high likelihood that the decision to adopt the EDDIE intervention was cost-effective. When an alternate willingness to pay of $0 per QALY was adopted, a mean NMB of $2,506 (SD $2,799) was estimated with 85% of simulations remaining positive.

## Discussion

The 12 months following the commencement of the EDDIE intervention was associated with a 19% reduction in annual hospital admissions and a 31% reduction in the average length of stay per admission when compared to the previous 12 months. When outcomes were modelled in a cohort of 1,000 RACF residents the intervention was estimated to produce an additional 0.62 QALYs while saving $2.6 million to the health care system. After accounting for plausible uncertainty in the model, there was an 86% chance of the intervention being cost-effective when adopting a willingness to pay of $28,000 per QALY. When the willingness to pay for health benefits was assumed to be zero, there was still an 85% change of the intervention being cost-effective and in this case, cost-saving to the health care system.

This is the first economic evaluation of a hospital avoidance intervention in the aged care setting. Prospective data were collected on the number of subacute episodes managed within the RACF as well as on the implementation costs of the intervention, including staff time spent on training, stakeholder engagement and project management activities. This information may be valuable to other RACFs considering adopting a similar program.

The study was limited to a single RACF in a regional area, and it is therefore unknown how the results we have reported may translate to other settings. A further limitation was the nonrandomised nature of the study design which may have introduced selection bias. As the intervention and usual care cohorts encompassed non-static resident populations it was not feasible to summarise and control for resident characteristics across the pre and post intervention periods. The analysis would have been further strengthened by the collection of prospective utility data which may be more sensitive to changes in the overall quality of care provided within the RACF.

Our findings support the growing body of evidence to suggest that programs allowing for sub-acute care to be provided within the RACF setting improve both resident and health service outcomes. Previous Australian studies have evaluated hospital in the nursing home (HINH) programs or other hospital or emergency department (ED) led outreach services that assist with the assessment of deteriorating residents^26 38 39^. These evaluations have reported significant reductions in ED transfers and hospital admission rates, but did not assess cost-effectiveness. The EDDIE intervention was instead focussed on upskilling existing RACF nursing staff and empowering them to proactively detect and respond to early signs of resident deterioration. In this sense it takes a similar approach to the hospital avoidance program ‘Interventions to Reduce Acute Care Transfers’ (INTERACT II) developed in the United States and reported to have reduced hospital admissions by 17-24% across 24 nursing homes^24^. Length of stay and cost-effectiveness outcomes associated with the INTERACT II program have not been reported. However, the EDDIE intervention is unique in that it was developed in, and driven by, the aged care setting.

The 2011 Australian Productivity Commission inquiry into caring for older Australians identified people in RACFs as being marginalised in terms of access to and quality of appropriate medical care ^40^. It was identified that continuity of care for RACF residents with acute healthcare needs and access to information of available services to fulfil their care needs were suboptimal. As identified by Ardents and Howard in their 2010 systematic review, older people living in RACFs have characteristics that distinguish them from the broader elderly population^6^. Notably, they are chronically ill and dependent, and the priority for their medical care is disease management rather than curative. In this context there is the opportunity for professional, accredited nursing staff to deliver appropriate care within the RACF and in turn prevent unnecessary hospital admissions.

The results of this evaluation are encouraging and provides compelling evidence to support the effectiveness of sub-acute care delivered by nursing staff in the RACF setting. The provision of a simple decision aid and staff training is a low cost intervention that may improve the quality of care residents receive while simultaneously providing high value to health systems by reducing the morbidity and expense associated with hospital transfers and admissions. Further implementation and evaluation of the EDDIE intervention in other RACFs is warranted to build a stronger evidence base around its effectiveness and cost-effectiveness.

## Acknowledgements

We thank the staff, residents and families at the Yaralla Aged Care Facility for their support in the implementation of the EDDIE intervention.

