## Appendix 1 for "The effectiveness and cost effectiveness of a hospital avoidance program in a residential aged care facility"

Costs of EDDIE implementation

| Implementation costs | $AUD |
| --- | --- |
| Decision support tool – printing costs | 360 |
| Project staff time on implementation activities |  |
| *Training and development (67 hrs)* | *2793* |
| *Stakeholder engagement (12 hrs)* | *494* |
| *Project management and leadership (14 hrs)* | *648* |
| **Total** | **4,295** |
