## Appendix 2 for "The effectiveness and cost effectiveness of a hospital avoidance program in a residential aged care facility"

Density of the fitted Normal distribution for the annual number of hospital admissions: usual care period (orange dashed line) and intervention period (blue solid line)

Density

| Annual number of admissions | Post subacute program | Pre subacute program |
| --- | --- | --- |
| 0 | 2.99662e-14 | 3.85221e-19 |
| 1 | 1.28817e-13 | 2.05614e-18 |
| 2 | 5.33011e-13 | 1.05869e-17 |
| 3 | 2.12285e-12 | 5.25847e-17 |
| 4 | 8.13806e-12 | 2.51955e-16 |
| 5 | 3.00291e-11 | 1.16456e-15 |
| 6 | 1.06656e-10 | 5.19246e-15 |
| 7 | 3.64625e-10 | 2.23336e-14 |
| 8 | 1.19985e-09 | 9.26655e-14 |
| 9 | 3.80039e-09 | 3.70896e-13 |
| 10 | 1.15864e-08 | 1.43205e-12 |
| 11 | 3.40009e-08 | 5.33384e-12 |
| 12 | 9.60396e-08 | 1.91644e-11 |
| 13 | 2.61114e-07 | 6.64237e-11 |
| 14 | 6.8333e-07 | 2.22088e-10 |
| 15 | 1.72127e-06 | 7.16312e-10 |
| 16 | 4.1734e-06 | 2.2287e-09 |
| 17 | 9.73978e-06 | 6.68923e-09 |
| 18 | 2.1879e-05 | 1.93675e-08 |
| 19 | 4.73072e-05 | 5.40936e-08 |
| 20 | 9.84568e-05 | 1.45744e-07 |
| 21 | 0.000197235 | 3.788e-07 |
| 22 | 0.000380314 | 9.49738e-07 |
| 23 | 0.000705864 | 2.29705e-06 |
| 24 | 0.00126101 | 5.35934e-06 |
| 25 | 0.00216838 | 1.20622e-05 |
| 26 | 0.003589 | 2.61887e-05 |
| 27 | 0.00571782 | 5.485e-05 |
| 28 | 0.00876815 | 0.000110819 |
| 29 | 0.0129421 | 0.000215985 |
| 30 | 0.0183874 | 0.000406075 |
| 31 | 0.0251453 | 0.000736486 |
| 32 | 0.0330989 | 0.00128853 |
| 33 | 0.0419362 | 0.00217471 |
| 34 | 0.0511429 | 0.00354063 |
| 35 | 0.0600345 | 0.00556075 |
| 36 | 0.0678322 | 0.00842481 |
| 37 | 0.0737719 | 0.0123129 |
| 38 | 0.0772264 | 0.0173594 |
| 39 | 0.0778145 | 0.0236093 |
| 40 | 0.0754701 | 0.0309746 |
| 41 | 0.0704545 | 0.0392014 |
| 42 | 0.0633086 | 0.0478598 |
| 43 | 0.0547566 | 0.0563657 |
| 44 | 0.0455859 | 0.0640372 |
| 45 | 0.0365295 | 0.0701816 |
| 46 | 0.0281758 | 0.0741974 |
| 47 | 0.0209185 | 0.0756706 |
| 48 | 0.0149487 | 0.0744458 |
| 49 | 0.0102824 | 0.0706524 |
| 50 | 0.0068078 | 0.0646825 |
| 51 | 0.0043385 | 0.0571244 |
| 52 | 0.00266128 | 0.0486664 |
| 53 | 0.00157131 | 0.0399955 |
| 54 | 0.000893005 | 0.0317079 |
| 55 | 0.000488501 | 0.0242492 |
| 56 | 0.000257215 | 0.0178896 |
| 57 | 0.000130361 | 0.0127314 |
| 58 | 6.35942e-05 | 0.00874034 |
| 59 | 2.98612e-05 | 0.00578833 |
| 60 | 1.34964e-05 | 0.00369787 |
| 61 | 5.87148e-06 | 0.0022789 |
| 62 | 2.45865e-06 | 0.00135479 |
| 63 | 9.90985e-07 | 0.000776946 |
| 64 | 3.84464e-07 | 0.000429818 |
| 65 | 1.4357e-07 | 0.000229379 |
| 66 | 5.16053e-08 | 0.000118085 |
| 67 | 1.78543e-08 | 5.86422e-05 |
| 68 | 5.94581e-09 | 2.80931e-05 |
| 69 | 1.9059e-09 | 1.29826e-05 |
| 70 | 5.8804e-10 | 5.78762e-06 |
| 71 | 1.74636e-10 | 2.48892e-06 |
| 72 | 4.99209e-11 | 1.03251e-06 |
| 73 | 1.37357e-11 | 4.13194e-07 |
| 74 | 3.63778e-12 | 1.59509e-07 |
| 75 | 9.2735e-13 | 5.94008e-08 |
| 76 | 2.27547e-13 | 2.13389e-08 |
| 77 | 5.37424e-14 | 7.3948e-09 |
| 78 | 1.22175e-14 | 2.47203e-09 |
| 79 | 2.67343e-15 | 7.97178e-10 |
| 80 | 5.63086e-16 | 2.47988e-10 |
| 81 | 1.14156e-16 | 7.44183e-11 |
| 82 | 2.22764e-17 | 2.15428e-11 |
| 83 | 4.18419e-18 | 6.01589e-12 |
| 84 | 7.56478e-19 | 1.62058e-12 |
| 85 | 1.31644e-19 | 4.21129e-13 |
| 86 | 2.20509e-20 | 1.05568e-13 |
| 87 | 3.55525e-21 | 2.55285e-14 |
| 88 | 5.51739e-22 | 5.95513e-15 |
| 89 | 8.24172e-23 | 1.34008e-15 |
| 90 | 1.18501e-23 | 2.90901e-16 |
| 91 | 1.64e-24 | 6.09161e-17 |
| 92 | 2.18468e-25 | 1.23053e-17 |
| 93 | 2.80124e-26 | 2.39789e-18 |
| 94 | 3.45727e-27 | 4.50752e-19 |
| 95 | 4.10711e-28 | 8.17373e-20 |
| 96 | 4.69633e-29 | 1.42981e-20 |
| 97 | 5.16893e-30 | 2.41272e-21 |
| 98 | 5.47599e-31 | 3.92746e-22 |
| 99 | 5.58399e-32 | 6.16723e-23 |
| 100 | 5.48083e-33 | 9.34205e-24 |

Density of the fitted Gamma distribution for length of stay: usual care cohort (orange dashed line) and intervention cohort (blue solid line).

Density

| Average length of stay | Post EDDIE |
| --- | --- |
| 1.06923e-05 | 1.53523e-119 |
| 1.65024e-05 | 2.95234e-115 |
| 0.000753388 | 1.53981e-77 |
| 0.000889003 | 6.62325e-76 |
| 0.000977135 | 5.67494e-75 |
| 0.00110272 | 8.85653e-74 |
| 0.00130592 | 4.1335e-72 |
| 0.00200744 | 7.22824e-68 |
| 0.0020166 | 8.01571e-68 |
| 0.00204662 | 1.12138e-67 |
| 0.00242436 | 5.25926e-66 |
| 0.00351382 | 2.41128e-62 |
| 0.00443143 | 4.68416e-60 |
| 0.00459386 | 1.06084e-59 |
| 0.00507013 | 9.96313e-59 |
| 0.00626171 | 1.20115e-56 |
| 0.00670118 | 5.60093e-56 |
| 0.00793261 | 2.57595e-54 |
| 0.00807539 | 3.8613e-54 |
| 0.00871598 | 2.18244e-53 |
| 0.0108244 | 2.97212e-51 |
| 0.0120393 | 3.31592e-50 |
| 0.0121847 | 4.3528e-50 |
| 0.0133093 | 3.21999e-49 |
| 0.0133887 | 3.68477e-49 |
| 0.0147313 | 3.2131e-48 |
| 0.016698 | 5.49365e-47 |
| 0.0172933 | 1.21451e-46 |
| 0.0176717 | 1.98272e-46 |
| 0.0193289 | 1.50872e-45 |
| 0.020379 | 4.99624e-45 |
| 0.0215346 | 1.74057e-44 |
| 0.0219099 | 2.57296e-44 |
| 0.0222018 | 3.47116e-44 |
| 0.0229047 | 7.0256e-44 |
| 0.0251 | 5.56607e-43 |
| 0.0253231 | 6.79905e-43 |
| 0.0280637 | 6.93528e-42 |
| 0.0282735 | 8.20622e-42 |
| 0.0291218 | 1.60021e-41 |
| 0.0330244 | 2.73788e-40 |
| 0.0379666 | 6.36404e-39 |
| 0.0383581 | 8.02003e-39 |
| 0.0387485 | 1.00764e-38 |
| 0.039135 | 1.26033e-38 |
| 0.0401252 | 2.21334e-38 |
| 0.0410787 | 3.75739e-38 |
| 0.0417108 | 5.30047e-38 |
| 0.0418328 | 5.66083e-38 |
| 0.0442308 | 1.98649e-37 |
| 0.045124 | 3.11585e-37 |
| 0.0451481 | 3.15346e-37 |
| 0.0453595 | 3.50318e-37 |
| 0.0463288 | 5.63822e-37 |
| 0.0466634 | 6.62961e-37 |
| 0.0504941 | 3.91084e-36 |
| 0.0507326 | 4.34791e-36 |
| 0.051009 | 4.913e-36 |
| 0.0516535 | 6.51539e-36 |
| 0.0520601 | 7.77117e-36 |
| 0.0538183 | 1.63932e-35 |
| 0.0550096 | 2.68101e-35 |
| 0.0557356 | 3.59921e-35 |
| 0.0558338 | 3.74439e-35 |
| 0.0561522 | 4.25443e-35 |
| 0.0562994 | 4.51199e-35 |
| 0.059298 | 1.44656e-34 |
| 0.059547 | 1.58918e-34 |
| 0.0600982 | 1.95425e-34 |
| 0.060237 | 2.05804e-34 |
| 0.0617623 | 3.60668e-34 |
| 0.0633297 | 6.32721e-34 |
| 0.0645028 | 9.54829e-34 |
| 0.065027 | 1.1448e-33 |
| 0.0659325 | 1.5608e-33 |
| 0.0681087 | 3.231e-33 |
| 0.0691504 | 4.53936e-33 |
| 0.0719729 | 1.11171e-32 |
| 0.0723208 | 1.23838e-32 |
| 0.072803 | 1.43697e-32 |
| 0.0761564 | 3.93499e-32 |
| 0.0770264 | 5.07278e-32 |
| 0.0790463 | 9.0478e-32 |
| 0.0801076 | 1.21893e-31 |
| 0.0803657 | 1.30975e-31 |
| 0.0807178 | 1.4441e-31 |
| 0.0810806 | 1.59631e-31 |
| 0.081229 | 1.66284e-31 |
| 0.0823262 | 2.24387e-31 |
| 0.0867503 | 7.21823e-31 |
| 0.0869005 | 7.50231e-31 |
| 0.0869878 | 7.67239e-31 |
| 0.089705 | 1.52356e-30 |
| 0.0970991 | 8.89671e-30 |
| 0.0980271 | 1.09947e-29 |
| 0.0991348 | 1.41184e-29 |
| 0.104673 | 4.72917e-29 |
| 0.106237 | 6.57591e-29 |
| 0.107172 | 7.98773e-29 |
| 0.108529 | 1.05634e-28 |
| 0.109319 | 1.24092e-28 |
| 0.110234 | 1.49322e-28 |
| 0.110567 | 1.59668e-28 |
| 0.113118 | 2.64874e-28 |
| 0.113748 | 2.99579e-28 |
| 0.121091 | 1.19865e-27 |
| 0.122627 | 1.58428e-27 |
| 0.129034 | 4.8888e-27 |
| 0.130724 | 6.51707e-27 |
| 0.137888 | 2.11709e-26 |
| 0.140309 | 3.10784e-26 |
| 0.143631 | 5.20597e-26 |
| 0.144644 | 6.07859e-26 |
| 0.144874 | 6.29449e-26 |
| 0.148032 | 1.01215e-25 |
| 0.148654 | 1.10993e-25 |
| 0.149787 | 1.31181e-25 |
| 0.152108 | 1.83994e-25 |
| 0.153598 | 2.27999e-25 |
| 0.162069 | 7.41481e-25 |
| 0.167279 | 1.48431e-24 |
| 0.168548 | 1.75183e-24 |
| 0.169232 | 1.91424e-24 |
| 0.170161 | 2.15838e-24 |
| 0.171759 | 2.6486e-24 |
| 0.174533 | 3.76136e-24 |
| 0.174592 | 3.7897e-24 |
| 0.175058 | 4.01716e-24 |
| 0.17527 | 4.12498e-24 |
| 0.176041 | 4.54059e-24 |
| 0.178389 | 6.06772e-24 |
| 0.182649 | 1.01637e-23 |
| 0.185771 | 1.47168e-23 |
| 0.18926 | 2.20895e-23 |
| 0.19067 | 2.59721e-23 |
| 0.191475 | 2.84715e-23 |
| 0.195808 | 4.63687e-23 |
| 0.196974 | 5.27695e-23 |
| 0.196989 | 5.28569e-23 |
| 0.198981 | 6.58037e-23 |
| 0.201177 | 8.35586e-23 |
| 0.20137 | 8.53148e-23 |
| 0.206735 | 1.51127e-22 |
| 0.215003 | 3.54093e-22 |
| 0.226489 | 1.09355e-21 |
| 0.230969 | 1.67028e-21 |
| 0.239664 | 3.70979e-21 |
| 0.241694 | 4.44991e-21 |
| 0.242523 | 4.7909e-21 |
| 0.245238 | 6.09015e-21 |
| 0.248255 | 7.92571e-21 |
| 0.251364 | 1.036e-20 |
| 0.253096 | 1.20089e-20 |
| 0.253419 | 1.23428e-20 |
| 0.254024 | 1.29927e-20 |
| 0.260536 | 2.23834e-20 |
| 0.268831 | 4.3848e-20 |
| 0.272555 | 5.8875e-20 |
| 0.273778 | 6.47978e-20 |
| 0.276141 | 7.78883e-20 |
| 0.280776 | 1.11191e-19 |
| 0.281714 | 1.19405e-19 |
| 0.283937 | 1.41239e-19 |
| 0.284532 | 1.47692e-19 |
| 0.286155 | 1.66771e-19 |
| 0.286193 | 1.67241e-19 |
| 0.288334 | 1.96075e-19 |
| 0.289517 | 2.13978e-19 |
| 0.292782 | 2.71796e-19 |
| 0.294724 | 3.12907e-19 |
| 0.294975 | 3.18637e-19 |
| 0.297445 | 3.80568e-19 |
| 0.300854 | 4.85063e-19 |
| 0.309144 | 8.64478e-19 |
| 0.310062 | 9.20658e-19 |
| 0.311505 | 1.01604e-18 |
| 0.313271 | 1.14552e-18 |
| 0.322364 | 2.10092e-18 |
| 0.326083 | 2.67822e-18 |
| 0.327928 | 3.01782e-18 |
| 0.330423 | 3.5423e-18 |
| 0.333013 | 4.17753e-18 |
| 0.34035 | 6.61719e-18 |
| 0.347208 | 1.00748e-17 |
| 0.354264 | 1.53834e-17 |
| 0.355927 | 1.69749e-17 |
| 0.35844 | 1.96785e-17 |
| 0.361346 | 2.33138e-17 |
| 0.36561 | 2.98184e-17 |
| 0.366362 | 3.11309e-17 |
| 0.371673 | 4.20857e-17 |
| 0.375319 | 5.16257e-17 |
| 0.376269 | 5.44268e-17 |
| 0.385558 | 9.05943e-17 |
| 0.386896 | 9.73872e-17 |
| 0.395669 | 1.55401e-16 |
| 0.397547 | 1.715e-16 |
| 0.407843 | 2.91801e-16 |
| 0.422858 | 6.1735e-16 |
| 0.425901 | 7.1604e-16 |

| Average length of stay | Pre EDDIE |
| --- | --- |
| 7.32493e-05 | 1.04829e-139 |
| 0.000279743 | 5.40622e-122 |
| 0.00064349 | 5.54155e-111 |
| 0.000823962 | 1.02596e-107 |
| 0.00202048 | 7.33482e-96 |
| 0.00857855 | 9.24898e-77 |
| 0.00879353 | 1.96269e-76 |
| 0.00923943 | 8.82806e-76 |
| 0.0106703 | 7.0224e-74 |
| 0.020834 | 4.70085e-65 |
| 0.0284187 | 5.77789e-61 |
| 0.0305813 | 5.33543e-60 |
| 0.0328936 | 4.85661e-59 |
| 0.0346402 | 2.32782e-58 |
| 0.0411747 | 4.35472e-56 |
| 0.0432211 | 1.88931e-55 |
| 0.0435543 | 2.38326e-55 |
| 0.0526174 | 7.22752e-53 |
| 0.0543862 | 1.96214e-52 |
| 0.0585962 | 1.8639e-51 |
| 0.0595757 | 3.07407e-51 |
| 0.0615192 | 8.09835e-51 |
| 0.06532 | 4.93989e-50 |
| 0.0675918 | 1.38477e-49 |
| 0.0676511 | 1.42188e-49 |
| 0.0688032 | 2.36546e-49 |
| 0.0753152 | 3.60651e-48 |
| 0.0810284 | 3.25752e-47 |
| 0.0870361 | 2.7994e-46 |
| 0.0924807 | 1.73352e-45 |
| 0.102351 | 3.63817e-44 |
| 0.103906 | 5.71863e-44 |
| 0.114061 | 9.35097e-43 |
| 0.120286 | 4.58756e-42 |
| 0.131665 | 6.84131e-41 |
| 0.138598 | 3.16534e-40 |
| 0.138749 | 3.27003e-40 |
| 0.143479 | 8.88751e-40 |
| 0.145716 | 1.40965e-39 |
| 0.146337 | 1.60017e-39 |
| 0.15202 | 4.97915e-39 |
| 0.156887 | 1.27217e-38 |
| 0.160994 | 2.7446e-38 |
| 0.161291 | 2.89895e-38 |
| 0.162272 | 3.47208e-38 |
| 0.163752 | 4.54845e-38 |
| 0.164507 | 5.21498e-38 |
| 0.168753 | 1.11194e-37 |
| 0.169795 | 1.33515e-37 |
| 0.171489 | 1.79304e-37 |
| 0.174367 | 2.93914e-37 |
| 0.179514 | 6.96809e-37 |
| 0.180952 | 8.82742e-37 |
| 0.181982 | 1.04456e-36 |
| 0.186119 | 2.03406e-36 |
| 0.191754 | 4.9213e-36 |
| 0.192261 | 5.32089e-36 |
| 0.195484 | 8.70402e-36 |
| 0.195523 | 8.7558e-36 |
| 0.198719 | 1.4145e-35 |
| 0.200008 | 1.71258e-35 |
| 0.211222 | 8.58568e-35 |
| 0.22124 | 3.3694e-34 |
| 0.223891 | 4.78634e-34 |
| 0.225315 | 5.76972e-34 |
| 0.226934 | 7.12427e-34 |
| 0.227017 | 7.20178e-34 |
| 0.234272 | 1.81834e-33 |
| 0.239092 | 3.31019e-33 |
| 0.24038 | 3.87678e-33 |
| 0.244343 | 6.2687e-33 |
| 0.248698 | 1.05331e-32 |
| 0.256035 | 2.47242e-32 |
| 0.267971 | 9.39829e-32 |
| 0.27219 | 1.48466e-31 |
| 0.285424 | 5.94851e-31 |
| 0.286323 | 6.52062e-31 |
| 0.291975 | 1.15365e-30 |
| 0.297992 | 2.09156e-30 |
| 0.324448 | 2.4845e-29 |
| 0.324641 | 2.52774e-29 |
| 0.329937 | 4.04334e-29 |
| 0.332763 | 5.17802e-29 |
| 0.339829 | 9.52133e-29 |
| 0.34166 | 1.11246e-28 |
| 0.359813 | 4.97242e-28 |
| 0.374657 | 1.59647e-27 |
| 0.379408 | 2.29523e-27 |
| 0.38686 | 4.01765e-27 |
| 0.399219 | 9.91963e-27 |
| 0.403645 | 1.3613e-26 |
| 0.403734 | 1.36992e-26 |
| 0.40391 | 1.38717e-26 |
| 0.405045 | 1.50348e-26 |
| 0.419427 | 4.08665e-26 |
| 0.423986 | 5.5684e-26 |
| 0.428325 | 7.45094e-26 |
| 0.434808 | 1.14456e-25 |
| 0.438685 | 1.47476e-25 |
| 0.44195 | 1.82226e-25 |
| 0.445159 | 2.23985e-25 |
| 0.446345 | 2.41641e-25 |
| 0.448391 | 2.75304e-25 |
| 0.464002 | 7.29353e-25 |
| 0.47004 | 1.05327e-24 |
| 0.480403 | 1.956e-24 |
| 0.48073 | 1.99415e-24 |
| 0.487985 | 3.04913e-24 |
| 0.489537 | 3.33606e-24 |
| 0.512796 | 1.23948e-23 |
| 0.519692 | 1.80676e-23 |
| 0.531011 | 3.31516e-23 |
| 0.531269 | 3.36091e-23 |
| 0.531952 | 3.48452e-23 |
| 0.535663 | 4.23734e-23 |
| 0.541473 | 5.73887e-23 |
| 0.549194 | 8.54087e-23 |
| 0.552142 | 9.92522e-23 |
| 0.552802 | 1.02637e-22 |
| 0.560829 | 1.53755e-22 |
| 0.561087 | 1.55749e-22 |
| 0.56387 | 1.78906e-22 |
| 0.602348 | 1.13012e-21 |
| 0.604793 | 1.26496e-21 |
| 0.609533 | 1.57178e-21 |
| 0.611679 | 1.73314e-21 |
| 0.615001 | 2.01469e-21 |
| 0.62661 | 3.38569e-21 |
| 0.632435 | 4.37569e-21 |
| 0.637621 | 5.48646e-21 |
| 0.64647 | 8.03332e-21 |
| 0.654039 | 1.10813e-20 |
| 0.661816 | 1.53561e-20 |
| 0.668431 | 2.01998e-20 |
| 0.676676 | 2.83108e-20 |
| 0.677661 | 2.94675e-20 |
| 0.679471 | 3.17104e-20 |
| 0.68471 | 3.91677e-20 |
| 0.689004 | 4.65072e-20 |
| 0.69013 | 4.86395e-20 |
| 0.719792 | 1.5406e-19 |
| 0.722778 | 1.72516e-19 |
| 0.734662 | 2.69236e-19 |
| 0.752882 | 5.24616e-19 |
| 0.760601 | 6.92166e-19 |
| 0.762663 | 7.44954e-19 |
| 0.764556 | 7.96814e-19 |
| 0.764972 | 8.08669e-19 |
| 0.769859 | 9.61145e-19 |
| 0.793921 | 2.21077e-18 |
| 0.797096 | 2.46234e-18 |
| 0.800124 | 2.72771e-18 |
| 0.812018 | 4.06041e-18 |
| 0.81433 | 4.38349e-18 |
| 0.825956 | 6.41841e-18 |
| 0.849147 | 1.34903e-17 |
| 0.858221 | 1.79275e-17 |
| 0.873684 | 2.88801e-17 |
| 0.875389 | 3.04216e-17 |
| 0.885408 | 4.11946e-17 |
| 0.887493 | 4.38553e-17 |
| 0.89392 | 5.31343e-17 |
| 0.904995 | 7.36867e-17 |
| 0.911253 | 8.84626e-17 |
| 0.911429 | 8.89153e-17 |
| 0.912074 | 9.05972e-17 |
| 0.912086 | 9.06289e-17 |
| 0.916707 | 1.03612e-16 |
| 0.928031 | 1.43382e-16 |
| 0.930507 | 1.5384e-16 |
| 0.946122 | 2.38678e-16 |
| 0.949594 | 2.62859e-16 |
| 0.960848 | 3.58413e-16 |
| 0.962688 | 3.769e-16 |
| 0.969777 | 4.57002e-16 |
| 0.973178 | 5.00996e-16 |
| 0.973916 | 5.11048e-16 |
| 0.975354 | 5.31217e-16 |
| 0.980778 | 6.14362e-16 |
| 0.987325 | 7.31307e-16 |
| 0.990406 | 7.93446e-16 |
| 1.00401 | 1.13324e-15 |
| 1.02231 | 1.81508e-15 |
| 1.03409 | 2.44463e-15 |
| 1.03813 | 2.70538e-15 |
| 1.04059 | 2.87685e-15 |
| 1.04224 | 2.99732e-15 |
| 1.04891 | 3.53642e-15 |
| 1.05747 | 4.36517e-15 |
| 1.06033 | 4.68037e-15 |
| 1.07036 | 5.97027e-15 |
| 1.08469 | 8.41265e-15 |
| 1.08903 | 9.32282e-15 |
| 1.08952 | 9.43108e-15 |
| 1.09407 | 1.05004e-14 |
| 1.11069 | 1.54656e-14 |
| 1.11562 | 1.73252e-14 |
| 1.13353 | 2.60407e-14 |
| 1.1394 | 2.97081e-14 |
| 1.14509 | 3.37369e-14 |
