## Appendix 3 for "The effectiveness and cost effectiveness of a hospital avoidance program in a residential aged care facility"

Distribution of Net Monetary Benefit across 1,000 Monte Carlo Simulations
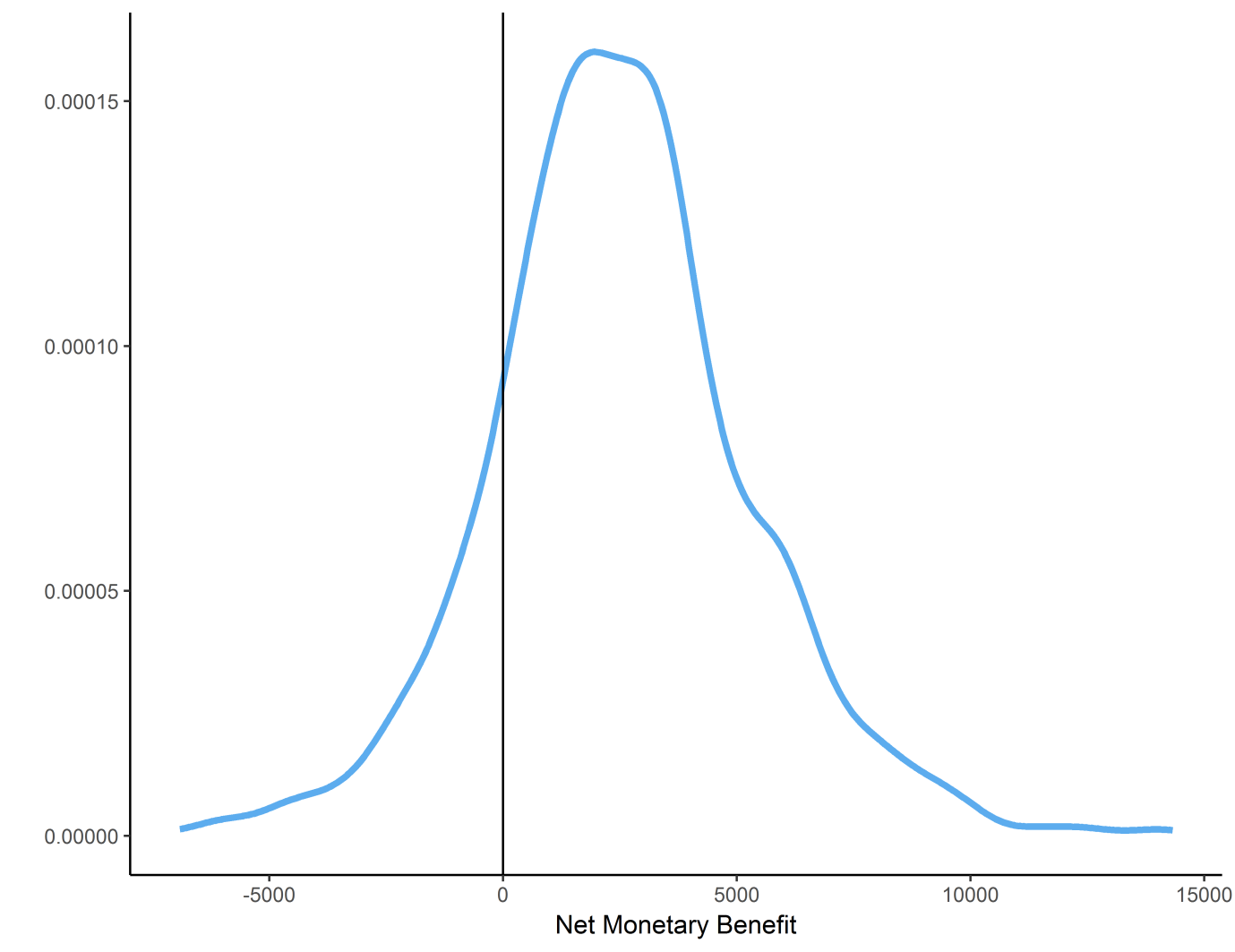

*NMB based on a willingness to pay of $28,000 per QALY
